# Multivalent dynamic colocalization of avian influenza polymerase and nucleoprotein by intrinsically disordered ANP32A reveals the molecular basis of human adaptation

**DOI:** 10.1101/2023.07.08.548184

**Authors:** Aldo R. Camacho-Zarco, Lefan Yu, Tim Krischuns, Selin Dedeoglu, Damien Maurin, Guillaume Bouvignies, Thibaut Crépin, Rob W.H. Ruigrok, Stephan Cusack, Nadia Naffakh, Martin Blackledge

## Abstract

Adaptation of avian influenza RNA polymerase (FluPol) to human cells requires mutations on the 627-NLS domains of the PB2 subunit. The E627K adaptive mutation compensates a 33-amino-acid deletion in the acidic intrinsically disordered domain of the host transcription regulator ANP32A, a deletion that restricts FluPol activity in mammalian cells. The function of ANP32A in the replication transcription complex and in particular its role in host restriction remain poorly understood. Here we characterise ternary complexes formed between ANP32A, FluPol and the viral nucleoprotein, NP, supporting the putative role of ANP32A in shuttling NP to the replicase complex. We demonstrate that while FluPol and NP can simultaneously bind distinct linear motifs on avian ANP32A, the deletion in the shorter human ANP32A blocks this mode of colocalization. NMR reveals that NP and human-adapted Pol, containing the E627K mutation, simultaneously bind the identical extended linear motif on human ANP32A, in an electrostatically driven, highly dynamic and multivalent ternary complex. This study reveals a probable molecular mechanism underlying host adaptation, whereby E627K, which enhances the basic surface of the 627 domain, is selected to confer the necessary multivalent properties to allow ANP32A to colocalize NP and FluPol in human cells.

## INTRODUCTION

Influenza A viruses (IAV) infect both mammals and aquatic birds, their natural hosts. Some highly pathogenic zoonotic avian strains, such as H5N1, can adapt to infect humans with high mortality, posing a serious pandemic threat.^1,2^ Genome replication in IAV is performed by the RNA-dependent RNA polymerase (FluPol), a heterotrimer, comprising three proteins, PA, PB1 and PB2 that assemble in the nucleus of the infected cell.^3^ The viral genome is made up of eight single-stranded, negative-sense RNA segments, each encapsidated by multiple viral nucleoproteins (NPs), and associated with FluPol to form ribonucleoprotein complexes (vRNPs). Replication requires two steps, first forming a complementary (cRNA), positive sense copy of the genome, that is used as a template to synthesize the viral RNA (vRNA) replicate. This process has been suggested to involve two FluPol trimers, one intrinsic to the vRNP, responsible for synthesis, and a second that controls encapsidation.^4–7^

Among other adaptations, mutations on the surface of FluPol are necessary in order to efficiently replicate in human cells. Adaptive mutations of avian IAV FluPol that are necessary for human infection are located to the C-terminal ‘627’ and ‘NLS’ domains of PB2.^8,9^ This region of PB2 is highly dynamic,^8,10^ with “open” and “closed” forms that interchange in a conformational equilibrium,^11^ a dynamic character that is retained in the assembled polymerase, suggesting a role in the viral replication cycle.^12,13^ Perhaps most remarkably, mutation of position 627 from E to K in avian PB2 rescues polymerase activity and viral replication in mammalian cells, spurring intense investigation into the molecular origin of this phenomenon.^14–17^ Barclay and co-workers identified members of the ANP32 family, host transcription factors comprising a Leucine-rich region (LRR), followed by a highly acidic intrinsically disordered domain (IDD),^18,19^ as responsible for this adaptation.^20^ The IDD exhibits an important difference across species, with a 33 amino acid deletion in human ANP32A (*hu*ANP32A) compared to avian ANP32A, consisting of an avian-unique hydrophobic hexapeptide followed by a repeat sequence of the preceding 27 amino acids (149-176). This avian-specific insertion alone was also shown to be sufficient to rescue avian-adapted H5N1 FluPol activity in mammalian cells, suggesting a mechanistic correlation between the E627K mutation and these differences in host ANP32A.^21–31^ Numerous studies have further identified the interaction of ANP32A, and the related ANP32B,^27–29^ with FluPol as critical to replication.^23,25,26^ We recently described the complexes formed between the 627-NLS domains of human- and avian-adapted FluPol (627(K)-NLS and 627(E)-NLS respectively) with *hu*ANP32A and chicken ANP32A (*ch*ANP32A),^32,33^ revealing that while a unique hexapeptide present in the avian IDD binds to 627(E)-NLS, the interaction is very different in human-adapted complex, where the E627K mutation completes a basic surface on the 627 domain,^9^ thereby allowing multivalent, transient interactions with a 50-amino acid stretch of the predominantly acidic IDD to stabilize the highly dynamic *hu*ANP32A:627(K)-NLS complex.

ANP32A is thought to be required primarily for viral genome replication,^21,26,34,35^ and it has recently been shown that 627E restricts vRNA but not cRNA synthesis in mammalian cells.^36^ Fodor and co-workers recently determined the structures of the complexes of *hu*ANP32A and *ch*ANP32A with FluPol from influenza C, bound to a 47-nucleotide vRNA.^37^ FluPol adopts an asymmetric dimeric form, comprising an active replicase, and a second FluPol, apparently acting as an encapsidase. In complex the folded domain of ANP32A bridges the two 627 domains.

Despite these advances, the role of ANP32A in influenza replication is not understood at the molecular level. Many single-stranded, negative-sense RNA viruses express a phosphoprotein (P), that chaperones the nucleoprotein (N),^38–41^ protecting against non-specific binding to cellular RNA prior to encapsidation, and controlling nucleoprotein concentration in the vicinity of FluPol, for example by forming membraneless organelles.^42–46^ Influenza does not express such a protein, raises the question of whether host ANP32A can compensate the absence of a viral adaptor protein, by performing similar functions. Recent biochemical, pull-down and mutagenesis studies have indeed demonstrated interaction of the disordered domain of ANP32A with NP, implicating the RNA-binding grooves on the surface of NP and the N-terminal half of the IDD of ANP32A as likely interaction sites.^47^

Here we describe the interaction of influenza NP with avian and human ANP32A at atomic resolution using NMR spectroscopy. We discover that the NP binding sites of ANP32A are located in the IDDs, and shifted by 33 amino acids, the length of the insertion, in *hu*ANP32A and *ch*ANP32A. The interaction is strongest in the avian form, where it is distinct from the 627(E)-NLS binding site, allowing both viral proteins to bind simultaneously on the same disordered chain, forming a ternary complex. In the shorter, human form, the two interaction sites collapse, so that the NP binding site precisely overlaps the 50 amino acid disordered stretch implicated in binding 627(K)-NLS. Despite this apparent overlap, NMR demonstrates the formation of a ternary complex, stabilized by multivalent interactions between *hu*ANP32A IDD and basic surfaces on both NP and 627(K)-NLS. On the basis of these results, we propose that ANP32A plays the role of an adaptor protein that can colocalize NP and FluPol, possibly for efficient encapsidation and stabilisation of newly synthesized RNA. Consideration of the intrinsic flexibility of the interacting domains of ANP32A and FluPol reveals that ANP32A can position NP in the putative RNA exit channel in both FluPol:*hu*ANP32A and FluPol:*ch*ANP32A complexes. The data demonstrate that while deletion of the 33 amino acid segment in *hu*ANP32A abrogates the ability to colocalise NP and FluPol using separate binding sites, by mutating E627 to K, FluPol acquires the ability to interact multivalently in an electrostatically driven ternary complex with the acidic disordered domain of *hu*ANP32A.

## RESULTS

### Isothermal titration calorimetry of NP interaction with avian and human ANP32A

Interaction between the monomeric mutant of H17N10 NP and *hu*ANP32A and *ch*ANP32A was initially investigated using ITC (figure S1). The interaction with *ch*ANP32A (120nM) is approximately one order of magnitude stronger than *hu*ANP32A (1.7μM). The enthalpic contribution is very similar in both cases, the difference lying predominantly in the increased entropic component. These affinities are notably stronger than those measured for the 627-NLS domains of FluPol which were estimated from NMR to lie in the 10s to 100s of micromolar range.^32^ A similar affinity of *hu*ANP32A for the R416A monomeric mutant nucleoprotein of the A/WSN/33(H1N1) was determined (1.8 μM - data not shown). The latter nucleoprotein was used for NMR experiments for reasons of stability.

### NMR reveals a shift in the NP interaction site between avian and human ANP32A

We investigated the interaction of ANP32A with the R416A monomeric mutant nucleoprotein of the A/WSN/33(H1N1) virus NP (hereon referred to as NP) using NMR spectroscopy (figure 1). Titration of NP into ^15^N labelled *ch*ANP32A results in localized reduction in intensity of peaks, with maximal reduction around ^210^EEYDEDAQVV^219^, but extending from 200-250. Direct interaction appears to be confined to the IDD, with negligible chemical shifts and uniform, weak broadening in the folded domain. No chemical shifts are observed in the centre of the binding region, with almost complete disappearance of peaks in this region at 1:0.5 admixture of NP, in agreement with the tight binding observed by ITC. Intensities in this region report on the residual free *ch*ANP32A in slow exchange between free and bound peaks, with bound peaks not visible due to the high molecular mass of the complex. Small shifts (e.g. 230E, 233E, 237S or 247Y) are observed in the peripheral regions associated with peaks in intermediate or fast exchange between free and bound protein.

**Figure 1.**
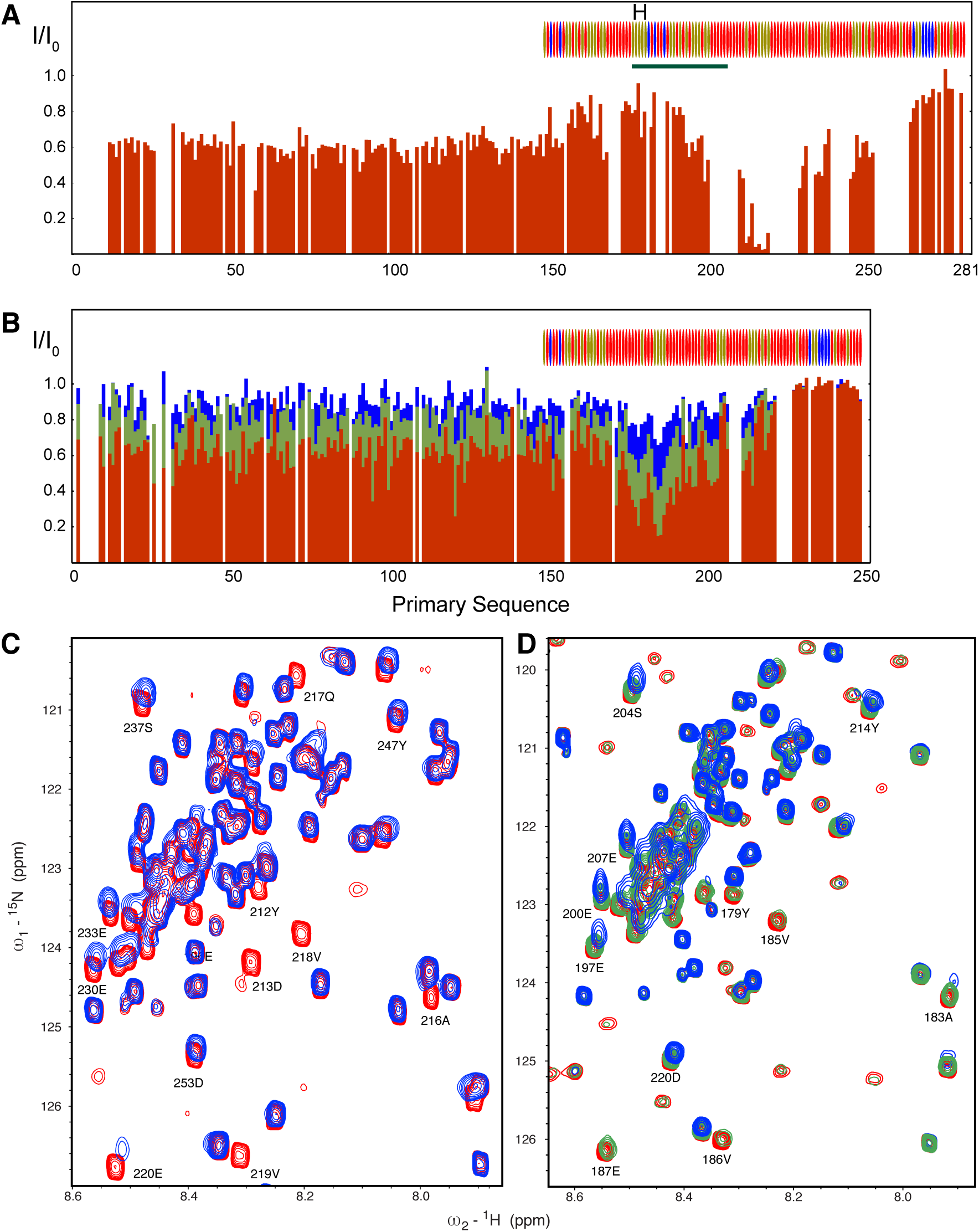
NP binds *ch* and *hu*ANP32A in the presence of increasing NP. (A) Intensities ratios of free and bound *ch*ANP32A. NP was added to 200 μM of ^15^N *ch*ANP32A at a 1:0.5 ratio (BEST-TROSY at 700 MHz). Bars show intensity ratio with respect to the free spectrum. The amino acid distribution of the IDD of *ch*ANP32A and *hu*ANP32A is shown above plots A and B respectively. Red = Asp/Glu, blue = Arg/Lys, beige = polar or hydrophobic residues. The avian IDD is 33 amino acids longer, comprising a hydrophobic hexapeptide (H) followed by a 27 amino acid repeat sequence (182-188) replicating the sequence (149-175). (B) Intensities ratios of free and bound *hu*ANP32A. NP was added to 100 μM ^15^N *hu*ANP32A at 1:0.05 (blue), 1:0.1 (green) and 1:0.2 (red/brown) ratio (BEST-TROSY at 850 MHz). Bars show intensity ratio with respect to the free spectrum. (C) BEST-TROSY (700 MHz) of free 200 μM of ^15^N *ch*ANP32A, (red) and at a 1:0.5 molar ratio of NP (blue). (D) BEST-TROSY (850MHz) of free ^15^N *hu*ANP32A at 100 μM (red), and at a 1:0.2 (green) and 1:0.5 (blue) molar ratio of NP.

Titration of NP into ^15^N labelled *hu*ANP32A results in a similar intensity profile, in this case shifted approximately 33 residues downstream, stretching from 169-221 and with the largest effects in the region ^177^EEYDEDAQVV^186^, a sequence that is identical to 210-219 in *ch*ANP32A. Small chemical shifts are observed as a function of admixture, accompanied by significant line broadening, suggesting intermediate exchange on the NMR timescale. Signal from the folded domain is further decreased compared to *ch*ANP32A interaction, possibly due to closer proximity of the binding site to the folded domain.

### NP binds ANP32A via the RNA binding pocket

In order to investigate the possible binding interface on the surface of NP, we titrated poly-GC RNA into the *hu*ANP32A:NP complex. This results in return of the intensity of the broadened ANP32A residues to the intensity of the free form, indicating that ANP32A and RNA binding is competitive, and implying that ANP32A binds in the RNA binding groove of NP (figure 2). Similar conclusions have been drawn from recent studies using pull-down assays of mutated NP with *hu*ANP32A, *hu*ANP32B and *ch*ANP32A.^47^ We note that a linear motif present in the ANP32A binding interface (EEYD) shares sequence identity with the conserved C-terminal peptide that binds on the opposite side of the RNA binding groove of NP, again supporting the suggestion that the negatively charged IDD of ANP32A binds the RNA binding groove of NP.^48,49^

**Figure 2.**
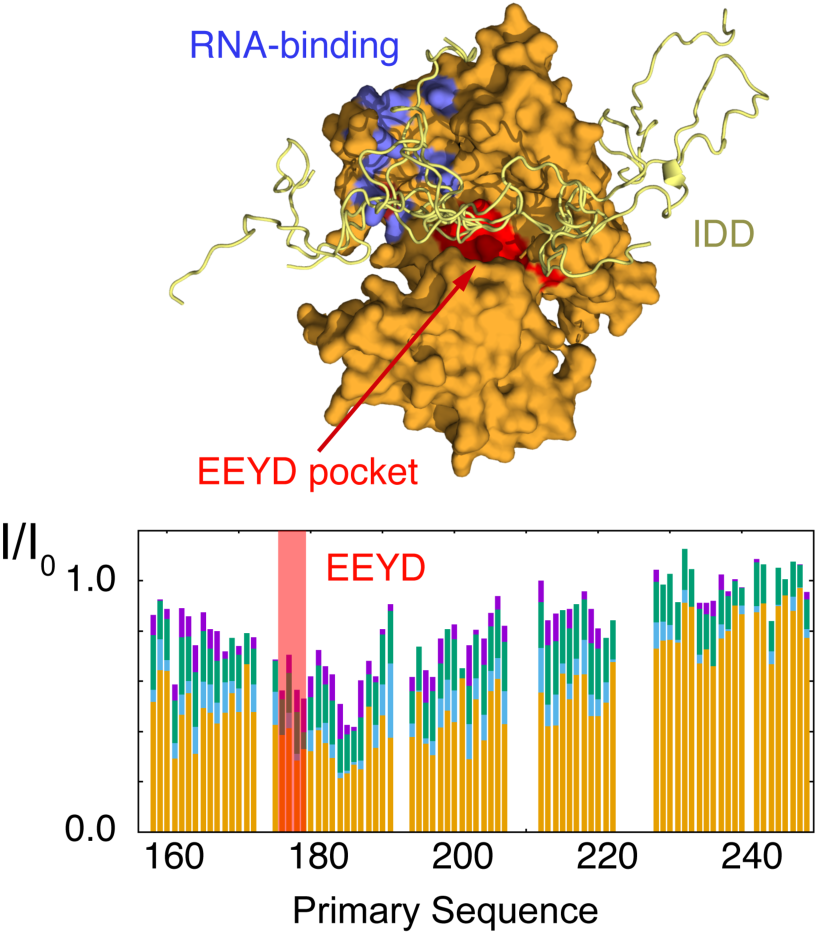
ANP32A IDD appears to bind in the RNA-binding pocket of NP. Histogram shows intensity recovery broadened resonances of ^15^N *hu*ANP32A (25μM) bound to H1N1 NP (1:1) upon addition of RNA. Yellow - *hu*ANP32A:NP (1:1), blue - 200% RNA poly UC 16mer, green - 200% RNA, purple – 800% RNA. Cartoon - Model of the IDD of *hu*ANP32A bound to NP; in blue the RNA binding grooves identified by Tang et al,^49^ red – the binding pocket locating the EEYD C-terminal peptide of NP.

### *Ch*ANP32A simultaneously binds NP and *av*627-NLS via distinct binding sites

In order to determine whether *ch*ANP32A can bind both NP and 627(E)-NLS, we titrated 627(E)-NLS into the *ch*ANP32A:NP complex. The affinity of the complex between *ch*ANP32A IDD is sufficiently tight that it is possible to study the 1:1 complex at a concentration of 100μM, essentially in the absence of the free forms of the proteins. Titration of 627(E)-NLS into this bimolecular complex resulted in almost identical shifts to those observed in the absence of NP (figure 3),^32^, with chemical shift perturbations (CSPs) again centered on the hexapeptide unique to *ch*ANP32A and part of the section that is deleted in *hu*ANP32A. This demonstrates that binding of 627(E)-NLS and NP to the two interaction sites on *ch*ANP32A, separated by 33 amino acids, are independent and can occur simultaneously, providing a molecular mechanism for colocalizing the two viral proteins via the host adaptor *ch*ANP32A. Spin relaxation (R_1ρ_) measured as a function of 627(E)-NLS admixture shows increasing relaxation in the peripheral regions (resonances from the interaction site are broadened beyond detection as expected for a complex of this size), further substantiating the observation of simultaneous binding of the two viral proteins and thereby formation of a ternary complex (figure 3).

**Figure 3.**
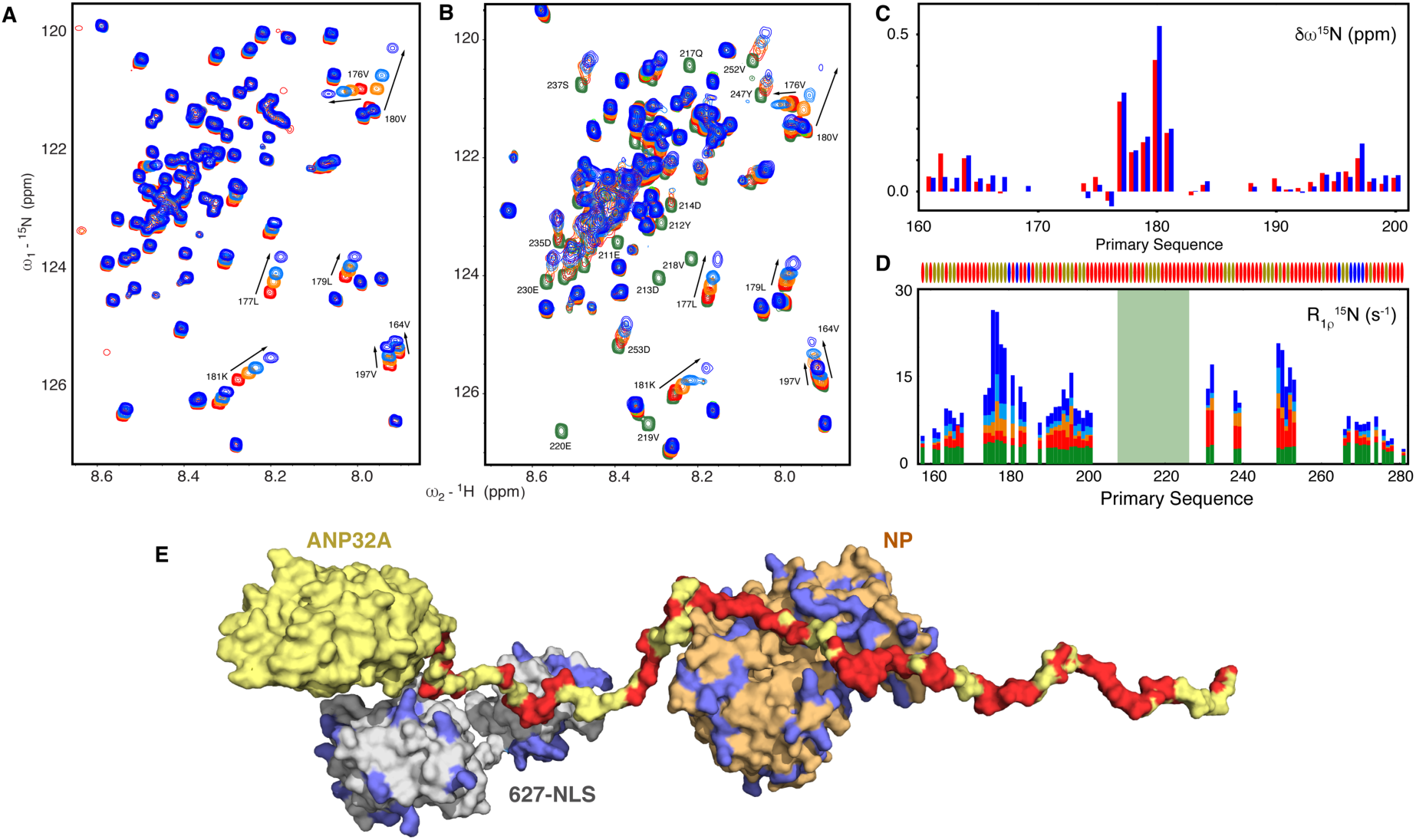
*ch*ANP32A IDD can simultaneously bind 627(E)-NLS and NP via two distinct binding sites. (A) Chemical shifts induced by titration of 627(E)-NLS into *ch*ANP32A IDD (25μM). Red – free *ch*ANP32A, orange – 100%, light blue – 200%, dark blue 400% 627(E)-NLS. These experiments were performed at 200mM NaCl.^32^ Spectra recorded at 850 MHz. (B) Chemical shifts induced by titration of 627(E)-NLS into *ch*ANP32A:NP 1:1 admixture (25μM). Green - free *ch*ANP32A IDD, Red – 1:1 *ch*ANP32A:NP admixture, orange – 100% 627(E)-NLS, light blue – 200% 627(E)-NLS, dark blue 400% 627(E)-NLS. Spectra recorded at 850 MHz. (C) ^15^N chemical shift differences between the 0% and 200% admixtures shown in A and B. Blue - *ch*ANP32A(IDD):627(E)-NLS, red - *ch*ANP32A:NP:627(E)-NLS (D) R_1ρ_ (850 MHz) of free *ch*ANP32A(IDD) (green); *ch*ANP32A(IDD):NP (1:1) (red); *ch*ANP32A(IDD):NP:627(E)-NLS (1:1:1) (orange); *ch*ANP32A(IDD):NP:627(E)-NLS (1:1:2) (light blue); *ch*ANP32A(IDD):NP (1:1:4) (dark blue). (E) Carton illustration demonstrating it is sterically possible for *ch*ANP32A (yellow) IDD (red shows positions of acidic residues) to simultaneously engage 627(E)-NLS (grey) and NP (orange) using the distinct binding sites characterized above (A-D). Positions of positively charged residues in NP and 627(E)-NLS are shown in blue.

### NMR exchange identifies the same interaction profile for NP and 627-NLS in human ANP32A

The suggestion of intermediate exchange in the *hu*ANP32A:NP interaction is verified by the measurement of ^15^N CPMG relaxation dispersion (figure 4 A,B), confirming the extensive interaction profile apparent from the intensity ratio (figure 1), affecting a stretch of more than 50 amino acids involved in the interaction. Data measured at two magnetic fields as a function of NP admixture (0, 5, 10 and 20%), are highly consistent, with the difference in ^15^N chemical shifts between free and bound states, over the entire region of *hu*ANP32A involved in the interaction agreeing very closely at all admixtures (figure S2). All data from each individual admixture can be fitted to the same 2-state process (figure S3), providing an estimation of the population of the bound state at each admixture, revealing site-specific affinities in the 50μM range. This is in contrast to the affinity measured for the entire molecular system from ITC (1.7μM), again indicating a strong component of multivalency, with interaction sites distributed over the 50 amino acid stretch contributing to the overall affinity. Remarkably, these chemical shift differences are essentially identical, both in magnitude and profile over the sequence, to those induced by interaction with human-adapted 627(K)-NLS (figure 4C),^32^ suggesting that the nature of the interaction is very similar in both cases, and that it involves the same long stretch of the IDD of *hu*ANP32A, forming polyvalent interactions with a strongly positively charged protein surface on NP or 627(K)-NLS.

**Figure 4.**
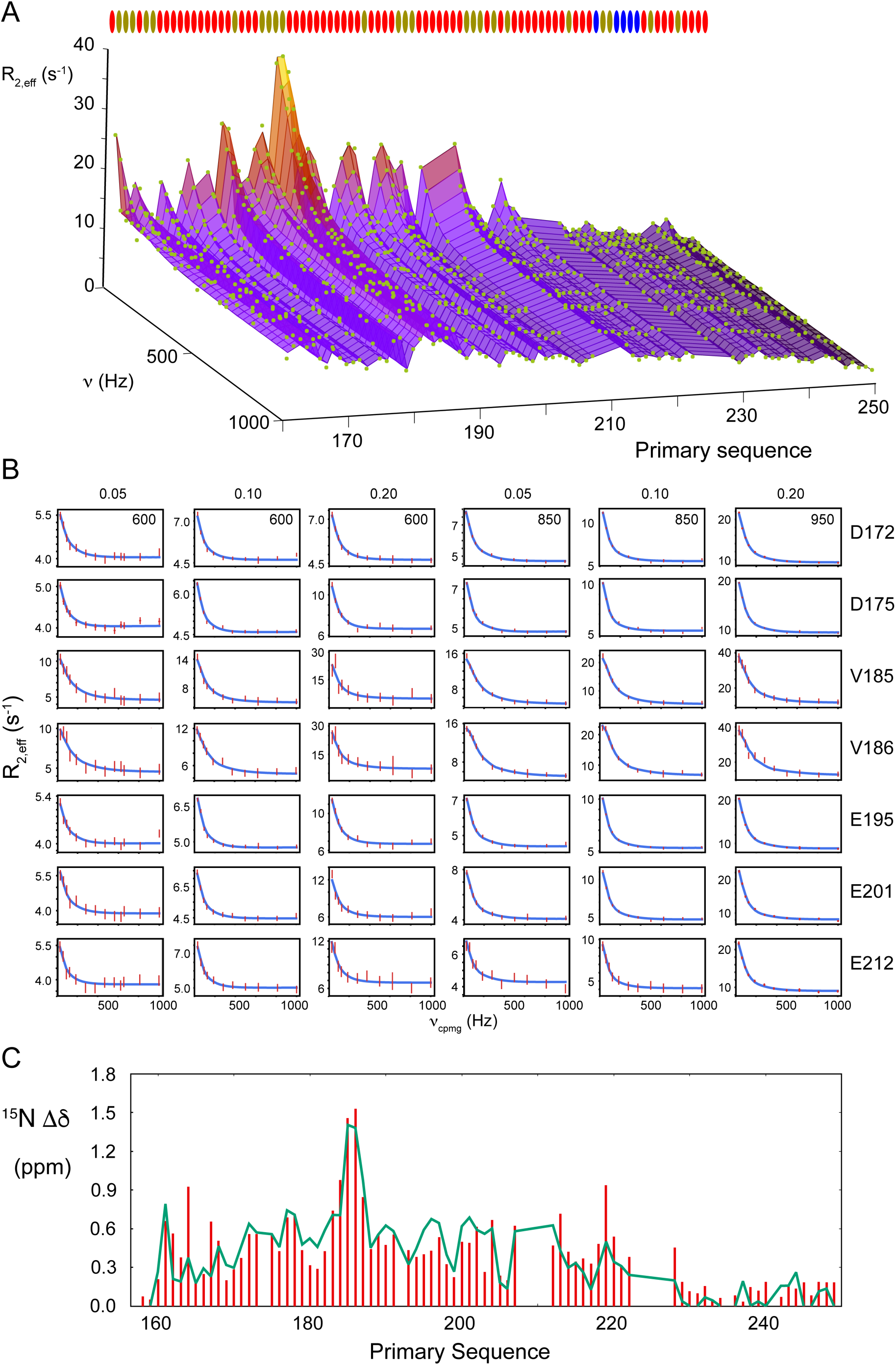
*hu*ANP32A binds NP and 627(K)-NLS via the IDD. (A) Relaxation dispersion measured throughout the IDD of *hu*ANP32A (950MHz). Data points are shown as green dots. The surface represents the simultaneous fit of all experimental data shown to a single 2-state exchange model (here the 20% admixture of NP is shown in 100μM *hu*ANP32A). The three-dimensional representation shows CPMG frequency, R_2,eff_ and primary sequence on three orthogonal axes. The colour code represents R_2,eff_. (B) CPMG relaxation dispersion profiles of 100μM ANP32A in the presence of 5%, 10% and 20% of NP showing selected residues from the region (172-212), measured at 600 MHz, 850 MHz, and 950 MHz as labelled. The entire data sets and fits are shown in figure S3. (C) Comparison of chemical shift differences between the free and bound form of *hu*ANP32A when it interacts with 627(K)-NLS (red) and NP (green). The former is derived from peak shifts from N-H HSQC spectra reporting on fast exchange, while the latter is extracted by fitting of CPMG relaxation dispersion data reporting on intermediate exchange.

### *hu*ANP32A simultaneously binds NP and 627(K)-NLS via the same binding site

The observation of simultaneous binding of NP and 627(E)-NLS to *ch*ANP32A, and the possible relevance of this ternary complex for colocalization of viral FluPol and NP in replication, poses the obvious question of whether *hu*ANP32A can colocalize NP and 627(K)-NLS. This is particularly intriguing given the apparently identical CSPs induced by both partners suggesting the identical interacting region along the disordered strand of *hu*ANP32A.

Addition of NP and 627(K)-NLS results in increase in transverse relaxation rates measured on *hu*ANP32A, as expected upon formation of the two binary complexes, with *hu*ANP32A being close to saturation in the 1:2 *hu*ANP32A: 627(K)-NLS complex. Addition of NP (10%) to this mixture results in an increase in transverse relaxation of *hu*ANP32A which is significantly and systematically higher than either of the two binary complexes (figure 5A). This suggests the existence of a cooperative binding mode, rather than competition between the bimolecular interactions of NP and 627(K)-NLS with *hu*ANP32A. While it is not possible on the basis of these data to estimate the correlation time of such a ternary complex, due to unknown stoichiometry and relative affinity, the most probable explanation for the higher relaxation rate is the simultaneous colocalization of all three proteins in a ternary complex. The same effect is observed for exchange-free relaxation measured using cross-correlated CSA-dipole cross-relaxation (η_xy_) (figure S4) indicating that this phenomenon is also not due to exchange contributions to R_2_. We note that no direct interaction was observed between NP and 627(K).

**Figure 5.**
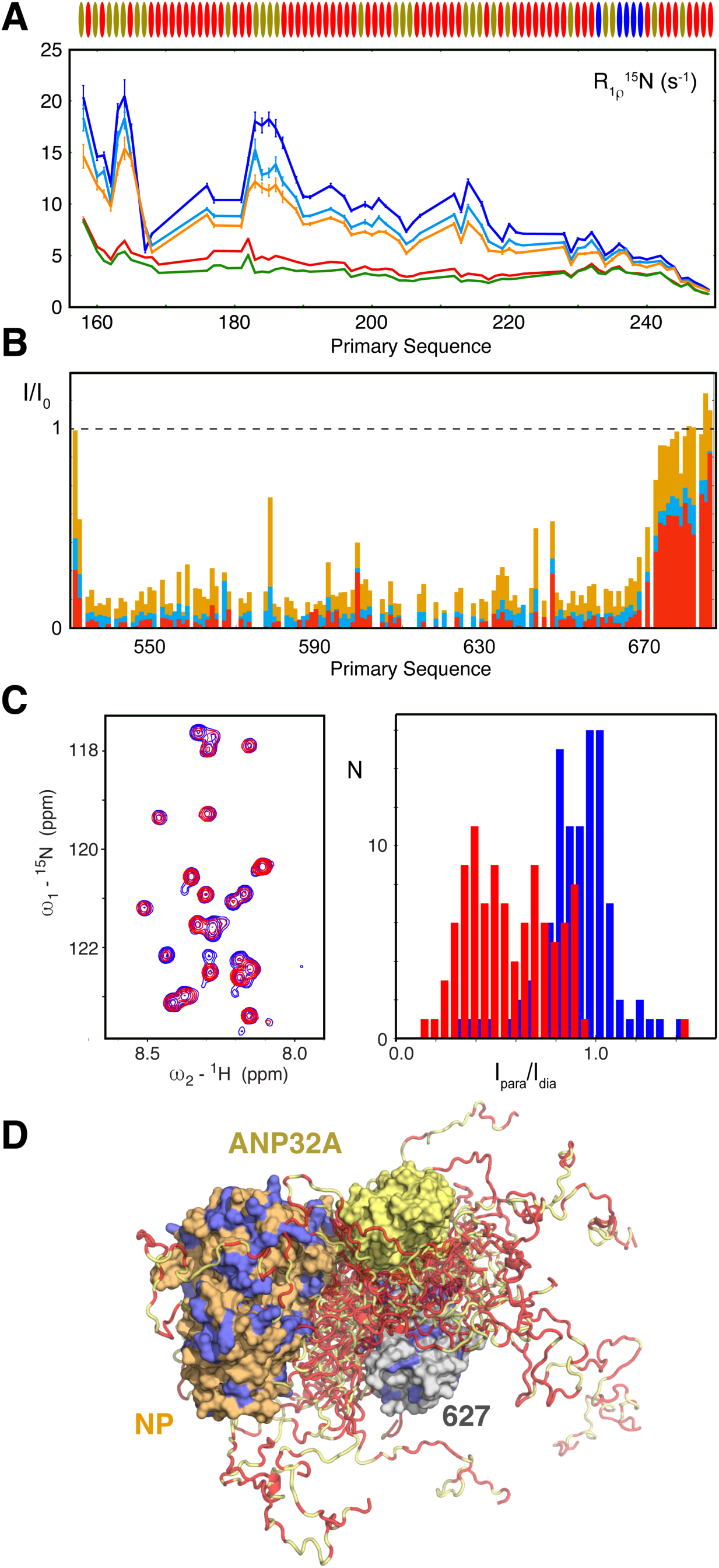
*hu*ANP32A forms a ternary complex with 627(K)-NLS and NP via the same binding interface. A) Residue-specific ^15^N R_1ρ_ of 100μM *hu*ANP32A in free form (green), in the presence of 10μM NP (red), in the presence of 100μM (orange) and 200μM (light blue) 627(K)-NLS, and in the presence of 10 μM NP as well as 200μM 627(K)-NLS (blue), demonstrating a clear increase in rotational correlation time as the ternary complex is formed. B) Intensity ratios of ^15^N-^1^H HSQC peaks of 100μM 627(K) relative to the free form, in the presence of 100μM *hu*ANP32A IDD (1:1:0 - orange), in the presence of 100μM *hu*ANP32A IDR and 100μM NP (1:1:1 - blue), and in the presence of 100μM *hu*ANP32A IDD and 200μM NP (1:1:2 - red). This again demonstrates that addition of NP does not compete with *hu*ANP32A binding. C) ^15^N-^1^H TROSY spectra and paramagnetic/diamagnetic intensity ratios of ^15^N labelled 100μM NP with 100μM 627(K) with a paramagnetic TEMPO label at residue 631, in the presence (red) and absence (blue) of 100μM *hu*ANP32A IDD. Distribution of extracted values of I_para_/I_dia_ for 100 resolved peaks is shown in histogram form (the peaks are not assigned). D) Multi-conformational cartoon representation of the interaction of *hu*ANP32A (yellow, with negatively charged amino acids shown in red) with both NP (orange, with positively charged amino acids shown in blue) and the 627(K) domain of FluPol (grey, with positively charged amino acids shown in blue).

Further evidence to support the existence of a ternary complex is provided by observing signals from ^15^N-labelled 627(K) in complex with unlabelled *hu*ANP32A. In the presence of *hu*ANP32A, the folded region of 627(K) uniformly lose intensity, presumably due to the increased correlation time of the complex (figure 5B). Upon addition of NP to this mixture, signals of 627(K) further lose intensity, rather than increasing as would be expected if 627(K) were released due to competitive binding of NP.

Finally, we observe approximately 100 ^15^N-^1^H TROSY cross peaks from the third component protein, NP. Although these are currently unassigned, they can be used to investigate the existence of contacts between NP and ANP32A and 627-NLS, by paramagnetically labelling 627(K) (at residue 631). We measured negligible line-broadening in NP spectra due to the presence of paramagnetic 627(K), confirming the lack of direct interaction with NP. However, when the experiment was repeated in the presence of both 627(K) and *hu*ANP32A (1:1:1), extensive paramagnetic broadening of numerous NP peaks is observed (figure 5C), confirming that a significant population of 627(K) and NP is only in close proximity when *hu*ANP32A is present. This verifies the observation that ANP32A is essential to colocalize the two viral factors and again directly demonstrates the existence of the ternary complex.

NMR relaxation and paramagnetic relaxation experiments from all three component proteins unambiguously confirm the formation of a ternary complex, mediated by the common extended interaction epitope on *hu*ANP32A. We previously identified the interaction between *hu*ANP32A and 627(K)-NLS as multivalent, on the basis of the very different site-specific affinities measured when observing *hu*ANP32A (600μM) or 627(K)-NLS (20μM), and the multiplicity of contacts along the extensive acidic binding interface on *hu*ANP32A interacting with the basic surface of 627(K)-NLS. Here we show that a similar phenomenon applies to the binding of NP, where relaxation dispersion identifies the identical extended binding surface on *hu*ANP32A, and a nearly 20-fold difference in affinity when observing the overall K_D_ and the local K_D_ determined for each site.

The ternary complex therefore exploits multivalent interactions between the acidic IDD of *hu*ANP32A and the basic surfaces of 627(K)-NLS and NP, allowing *hu*ANP32A to colocalize both partners. This novel interaction mode compensates the disappearance of the 627(E)-NLS binding site present in *ch*ANP32A, and provides a physical explanation for the crucial impact of the E627K mutation on adaptation.

### Distinct ternary NP:ANP32A:627-NLS complexes in human and avian adapted environments

The conformational sampling of 627-NLS and ANP32A have been previously described in detail in the case of both human-adapted and avian-adapted binary complexes.^32^ Here we investigate the steric feasibility of the proposed ternary complexes, incorporating NP in addition to the bound 627-NLS domains. Assuming the broad interaction interface comprises *ch*ANP32A ^210^EEYD^213^ bound in the RNA binding groove, it is clear that both viral partners can simultaneously bind *ch*ANP32A IDD (see cartoon in figure 3). Representing the human-adapted complex poses a more challenging problem, requiring the depiction of multivalent interactions involving both NP and 627(K)-NLS with *hu*ANP32A. The nature of the observed multivalency strongly suggests a highly dynamic complex, involving interaction sites distributed along the region between 170 and 220 binding and unbinding rapidly to both partners, thereby maintaining the colocalization of *hu*ANP32A with 627(K)-NLS or NP simultaneously in the ternary complex. In particular NP and 627-NLS need to be close enough to allow for paramagnetic broadening of residues on the surface of NP by the TEMPO label attached to residue 631 in 627(K) (see cartoon figure 5D). These putative multi-conformational models are presented to demonstrate the steric feasibility of such a ternary complex and to illustrate the rapid interchange between multivalent contact points that maintains ternary colocalization.

### Mutation studies confirm the multivalent nature of the *hu*ANP32A:NP interaction

The multivalent nature of the interaction is supported by the observation that mutation of the apparent interaction hotspots (as measured from the magnitude of the CSP), ^178^EEYDEDAQVV^185^ to ^178^AAAAEDAQGG^185^ do not abolish the interaction as observed on the remaining residues in the interaction interface. Relaxation dispersion is measured throughout the interaction interface (figure 6A), revealing similar chemical shift differences as in the wild-type for the non-mutated residues (figure 6B). Interestingly, the mutation increases relaxation rates in the upstream 200-220 region, suggesting a compensatory increase in the population of the remainder of the bound state upon mutation of these motifs (figure 6C). These data substantiate a model in which none of the individual linear motifs participating in the multivalent interaction are essential, and that each contributes to the overall affinity.

**Figure 6.**
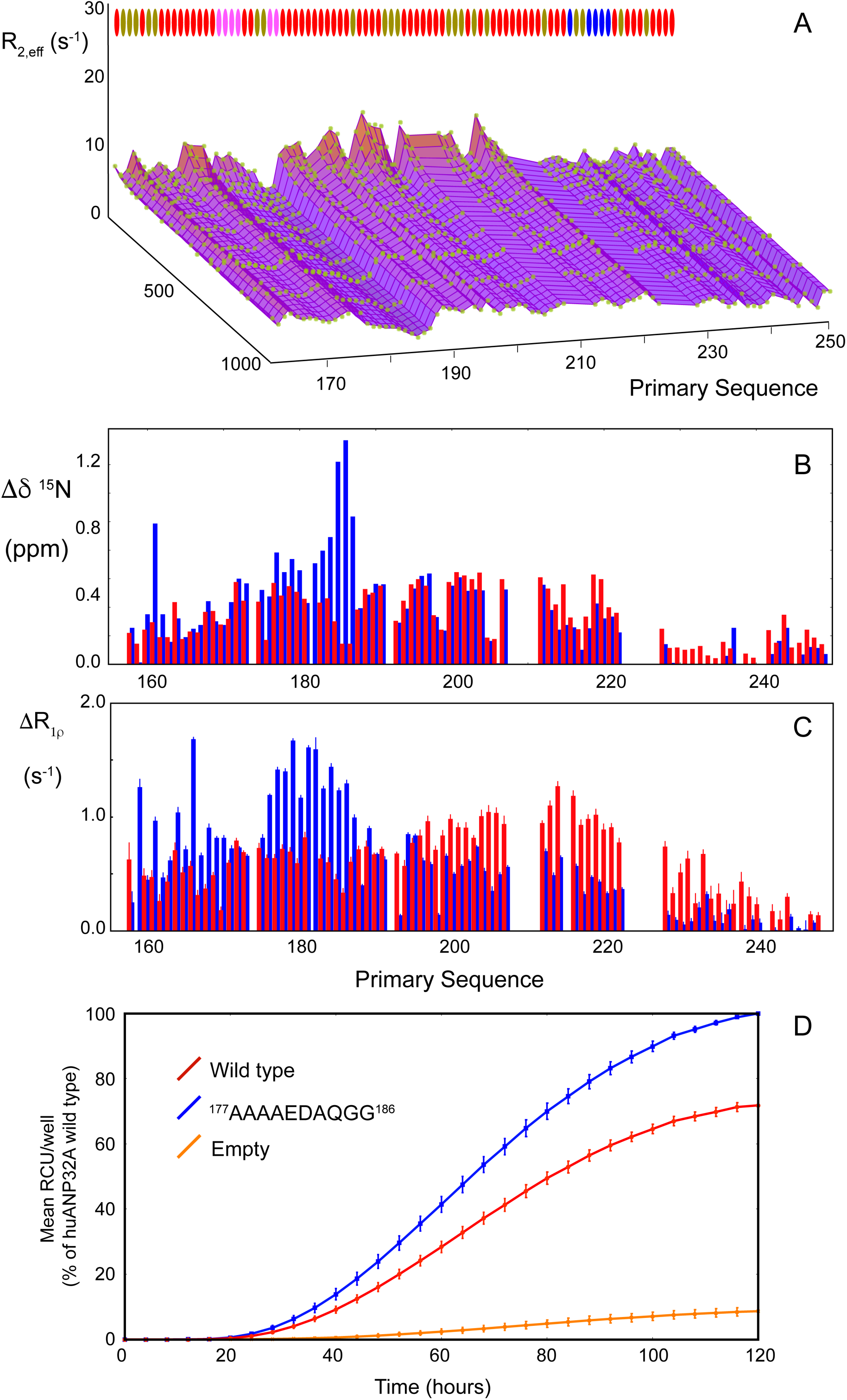
Relaxation and relaxation dispersion confirm the multivalent nature of the *hu*ANP32A:NP interaction. A) CPMG relaxation dispersion profile of 100μM *hu*ANP32A throughout the IDD (^177^AAAAEDAQGG^186^) in the presence of 10μM NP, measured at 850MHz. Data points are shown as green dots. The surface represents the simultaneous fit of all experimental data shown to a single 2-state exchange model. The three-dimensional representation shows CPMG frequency, R_2,eff_ and primary sequence on three orthogonal axes. The colour code represents R_2,eff_. B) Residue-specific chemical shift differences between the free and bound form of WT (blue bars) and *hu*ANP32A(^177^AAAAEDAQGG^186^) (red bars) (both 100 μM), in the presence of 10μM NP, as determined from relaxation dispersion measured at 600 and 850 MHz. All sites were fit simultaneously to a 2-state exchange model using ChemEx (p_B_=0.70±0.01, k_ex_ = 270±11s^-1^). C) Residue-specific ^15^N R_1ρ_ (700 MHz) differences of *hu*ANP32A in the presence and absence of 10μM NP; *hu*ANP32A WT (blue bars) and *hu*ANP32A(^177^AAAAEDAQGG^186^) (red bars). D) Cellular influenza polymerase activity in the absence of ANP32A/B (orange), and in the presence of *hu*ANP32A-wt (blue), and *hu*ANP32A ^177^AAAAEDAQGG^186^ (red), corresponding to the mutation investigated *in vitro*. A/WSN/1933 FluPol activity was measured by vRNP reconstitution in HEK-293T ANP32AB KO cells by transient transfection using a model vRNA encoding mCherry as well as either *hu*ANP32A wild-type (WT) or *hu*ANP32A mutants. Fluorescence signals were acquired every 4 h and are represented as RCU x µm²/Well ± SD of three experiments performed in triplicate.

Cellular assays were carried out to measure polymerase activities using ANP32A/B knock-out HEK293T cells complemented with transiently expressed wild-type or ^178^EEYDEDAQVV^185^ to ^178^AAAAEDAQGG^185^ mutated *hu*ANP32A, in a standard vRNP reconstitution assay using a viral model RNA encoding mCherry. Monitoring of mCherry fluorescence, measured as a function of time post-infection, demonstrates that while polymerase activity is maintained (consistent with studies *in vitro*) its overall efficiency is slightly reduced in the mutated form. This suggests, possibly not surprisingly, that the increased binding of the upstream region in the presence of these mutations is only partially compensating the defect in supporting polymerase activity.

### NP:ANP32A:627-NLS ternary complexes in the context of the full-length replicase/encapsidase

It is evident that the conformational space available for NP bound to ANP32A will be modified in the context of full-length FluPol. In this context we have modelled the conformation of a dimeric form of FluPol from IAV bound to ANP32A, using the recent structure of FluPol Influenza C.^37^ Following the proposition of Carrique et al, nascent RNA is threaded from the replicase FluPol active site, via the proposed RNA exit channel to the encapsidase FluPol where the 5’ RNA is tethered (figure 7A,B). The synthesized RNA is then solvent accessible in the exit channel. We note that the relative position of the folded domain of ANP32A and the 627 domain in both solution ensembles of 627-NLS and ANP32A is similar to the relative conformation of ANP32A and the encapsidase 627 domain, suggesting that this orientation is predominant in the minimal complex studied in solution. The conformational space available to the IDD in the context of this model was calculated using flexible-meccano,^50,51^ allowing us to investigate the possible sampling radius of ANP32A and the relevance of the proposed ternary complexes within this structural context. Assuming that NP binds 24 nucleotides of the newly synthesized RNA in the groove between FluPol_R_ and FluPol_E_, we impose a minimum distance of 40Å between the respective binding regions on *hu*ANP32A and *ch*ANP32A and RNA as it exits FluPol_R_. In the case of avian ANP32A, the NP-binding site can clearly reach the putative RNA exit channels, suggesting a plausible mechanism for locating NP to nascent RNA, while locating the 627-NLS binding site, 33 amino acids upstream, to the surface of 627-NLS (figure 7C).

**Figure 7.**
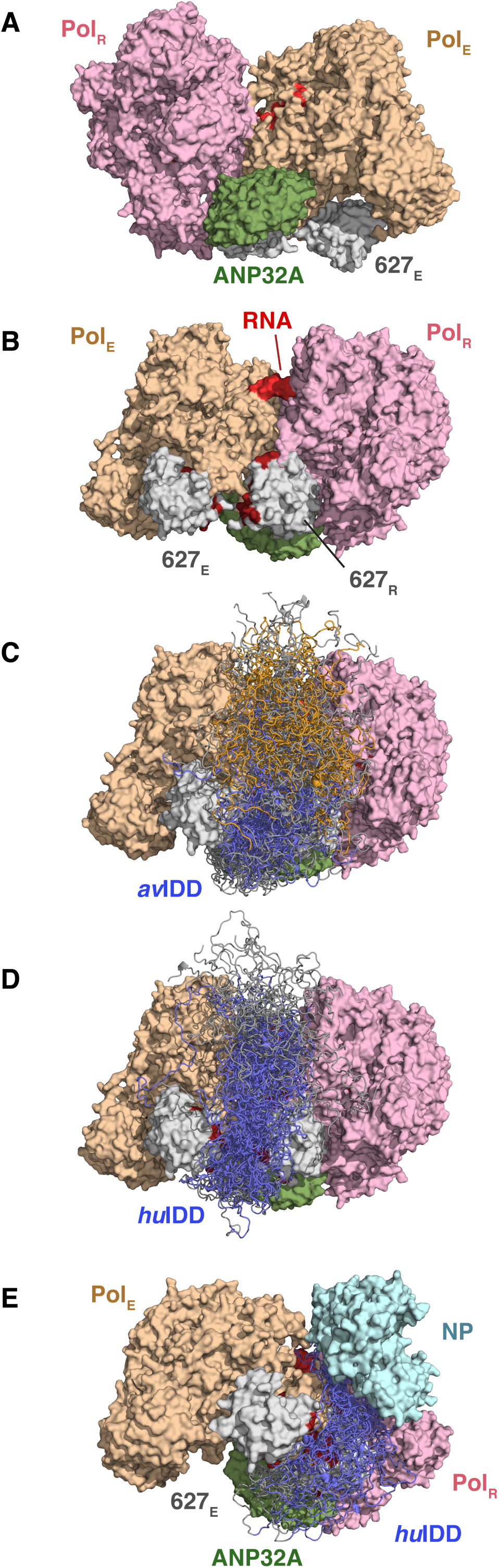
Modelling the ternary complexes of ANP32A in the context of the dimeric FluPol replicase. (A) The conformation of the replicase/encapsidase FluPol dimer was modelled on the basis of the Influenza C structure determined by Carrique et al. ^37^ Synthesized RNA was threaded from the replicase FluPol (pink), via the proposed RNA exit channel to the encapsidase FluPol (orange). The exit point of the synthesized RNA (red) is then solvent accessible, as seen in the rotated conformation (B). ANP32A folded domain (green) is located in the equivalent position as determined for Influenza C. The two 627 domains corresponding to the replicase (627_R_) and the encapsidase (627_E_) are shown in grey, with the ANP32A binding residues shown in dark red. (C) Conformational sampling of the disordered domain of *ch*ANP32A (*ch*IDD) was calculated using flexible-meccano. Conformers shown all fulfill the constraints that the 627(E) binding region (blue - the region 175-195, including the hexapeptide) is in the vicinity (<15Å) of one of the 627 domains, while the NP binding motif (orange, 205-225) is within 40Å of the exit point of the newly synthesized RNA (thereby allowing NP to bind the emergent RNA). This suggests a plausible mechanism for locating NP to nascent RNA, while locating the 627(E)-NLS binding site, 33 amino acids upstream, to the surface of 627(E)-NLS (figure 7C). (D) The equivalent *hu*ANP32A complex in the same context. The extended 627(K) and NP binding motif (blue), from 170:220, can sample conformations within 40Å of the RNA exit point as well as either 627_R_ or 627_E_. (E) NP was placed in the bottom of the exit channel, to illustrate the ability of the extended binding motif of *hu*ANP32A to locate NP within 40Å of the RNA exit point, while interacting with 627_R_/627_E_.

The equivalent *hu*ANP32A complex with FluPol can be modelled in a similar fashion. In this case we investigated whether the extended binding motif can sample conformations within 40Å of the RNA exit position, while interacting with the 627 domains, revealing that this is indeed the case, placing NP at the base of the broadly defined RNA exit channel (figure 7D). In order to accommodate this restraint, the ANP32A IDD is required to pass in the vicinity of the previously identified interaction surfaces of FluPol_R_ and FluPol_E_ 627 domains, comprising residues 587, 591, 627 and 631, all amino acids whose mutation has been associated with pandemic strains, and which show enhanced interaction in the *hu*ANP32A:627(K)-NLS complex compared to the *ch*ANP32A:627(E)-NLS complex.^32^ This also correlates closely with the observation of electron density between basic residues on the two closely positioned 627 domains in the influenza C dimer, density which was assigned to the acidic *hu*ANP32A IDD.^37^ In order to determine whether NP can be accommodated in this molecular assembly, we have positioned NP in the bottom of the RNA binding cleft (at 40Å from the initial RNA exit point), demonstrating that amino acids on the common binding epitope on *hu*ANP32A can intercalate between FluPol_R_ and FluPol_E_ 627-NLS domains and NP, and supporting the plausibility of such a pre-assembly configuration (figure 7E).

## DISCUSSION

The interaction between IAV FluPol and members of the host transcription factor ANP32 family has been shown to be essential for viral replication,^7,20,23,25–29^ although its function is not yet understood. In particular the presence of a 33-amino acid deletion in the human forms of the protein has been shown to drive adaptation of FluPol, via E627K or other amino acids on the surface of the 627-NLS domains, so that avian FluPol can function in human cells.^20^ The molecular function of ANP32 in viral replication is not known, although it has been shown to interact with both NP and FluPol, suggesting a role related to paramyxoviral phosphoproteins, essential viral cofactors that traffic nucleoproteins to the RNA replication-transcription complexes.^47,52–55^ Our recent study described the highly dynamic bimolecular complexes between human and avian ANP32A and human and avian adapted 627-NLS domains of FluPol, revealing distinct binding modes.^32^ The human complex exploits extensive multivalency, involving electrostatic interactions dispersed along the length of the highly acidic IDD and the 627 domain where the E627K mutation creates a continuous electrostatic surface. This kind of highly multivalent interaction involving IDPs has been observed throughout biology,^56–58^ for example between electrostatically complementary nuclear proteins,^59,60^ in the nuclear pore, involving short linear motifs,^61^ and playing a role in the stabilization of membraneless organelles.^62^ The avian complex on the other hand was found to be stabilized by interactions involving the avian-specific hexapeptide in the avian ANP32A IDR, binding to 627(E)-NLS, and no direct evidence for multivalency was detected.

Here we further investigate the role of ANP32A in IAV replication by describing the interactions of NP with avian and human ANP32A. NP interacts with the IDDs of ANP32A, via an extended binding site comprising a peptide motif (EEYD) that is also present in the C-terminal tail of NP, and that likely binds in the RNA binding cleft, as previously proposed.^47^ The NP interaction site on ANP32A is shifted by 33 amino acids between human and avian forms, which corresponds to the length of the avian-specific insertion. In the avian case, we show that NP interaction is fully compatible with binding of 627(E)-NLS, allowing the two viral proteins to colocalize simultaneously on the IDD, forming a ternary hub-like complex.

NP binding to *hu*ANP32A is an order of magnitude weaker (1.7μM) and in intermediate exchange on the NMR timescale, allowing us to precisely characterise the structure, kinetics and thermodynamics of the interaction using NMR exchange methods. This reveals that an extended stretch of more than 50 amino acids are involved in the interaction interface, with an effective site-specific affinity of around 50μM, indicating that similarly to the 627(K) domain, ANP32A also binds NP multivalently via multiple interaction sites along the chain, whose individual affinities are significantly weaker than the collective affinity for the entire protein. Most remarkably, the chemical shift differences induced by the interaction are essentially identical to those measured on the same protein upon interaction with the 627(K)-NLS domains of FluPol, both in amplitude and exact distribution along the primary sequence.

NMR relaxation of ANP32A measured in the presence of both partners reveals the formation of a ternary complex capable of significantly slowing the rotational correlation times. In addition, 627(K) resonances broadened due to addition of ANP32A are further broadened upon addition of NP, again indicating the absence of competitive binding, while broadening of backbone and sidechain resonances of NP, due to the addition of paramagnetically labelled 627(K), is only observed in the presence of ANP32A. These data, individually and collectively, demonstrate the existence of a ternary complex of 627(K)-NLS:*hu*ANP32A:NP, mediated by the same 50 amino-acid stretch of *hu*ANP32A. Remarkably, the IDD is clearly interacting multivalently with both folded viral partner proteins, so that fast on and off rates with respect to both proteins allow individual binding sites on the IDD to interact rapidly and transiently with the basic surfaces of the two partners thereby maintaining their dynamic colocalization. This observation reveals the importance of the previously observed multivalent *hu*ANP32A:627(K)-NLS interaction, unique to the human-adapted polymerase, and uncovers the molecular basis of the crucial E627K adaptive mutation, that confers the ability to interact multivalently and thereby enable ternary interaction, despite the absence of the two distinct binding sites present in *ch*ANP32A.

This remarkable multivalent nature of the interaction is further supported by investigation of the interaction of NP with *hu*ANP32A mutated at the apparent NP interaction hotspots (^178^EEYD^181^ and ^184^VV^185^). In this case although interaction at these sites is impacted, binding is enhanced for residues further along the binding region, as expected by increased avidity for these sites due to the absence of the mutated sites. Cellular assays conducted using ANP32A/B knock-out HEK293T cells expressing mutated ANP32A also demonstrate that while polymerase activity is affected when the ^178^EEYD^181^ and ^184^VV^185^ sites are mutated, it is not abolished. This also testifies to the resilience of this interaction mechanism to variations encountered in mammalian ANP32 IDDs.

Cryo-EM structures show that ANP32A binds to an asymmetric dimer of FluPol from influenza C, bridging across replication and encapsidation-competent FluPols.^37^ This dimeric interface was also found in crystal and cryo-EM structures in the absence of ANP32A, suggesting that the host factor is not essential to the structure, although it may stabilize the conformation, in particular because electron density was noted between basic residues on the two closely positioned 627 domains,^37^ and this density was assigned to the acidic *hu*ANP32A IDD.

Using this structure as a template, we modelled the replication/encapsidation dimer of IAV, threading an RNA strand from the replicase to the encapsidase. We used this model to explore the possible sampling behaviour of the *hu*ANP32A IDD in this molecular context, building the entire disordered chain in agreement with experimental NMR chemical shifts. Simple restraints, requiring simultaneous proximity of *hu*ANP32A residues in the interaction motif with the surface of 627(K) and placing NP at an appropriate distance to the RNA exit channel, demonstrate that the NP:627(K) multivalent interface on the IDD can indeed acts as an adaptor, colocalizing NP and 627 in the vicinity of newly synthesized RNA prior to encapsidation. We note that in order to fulfill these constraints, the extended binding motif of the *hu*ANP32A IDD must pass in the proximity of the interface between the replicase and encapsidase 627 domains, as previously suggested on the basis of electron density observed in this channel.^37^ Not surprisingly, *ch*ANP32A IDD can easily fulfill constraints that place the two interaction motifs in the vicinity of 627(E)-NLS and the putative RNA exit channel respectively.

FluPol is known to be extremely plastic in solution, as illustrated by the numerous high resolution structures,^3,5,12,13,63–65^ where PB2, and the 627-NLS domains in particular, have been shown to either change position relative to the core of FluPol,^13^ or to disappear entirely, indicating extensive sampling of conformational space on the surface of FluPol. It is possible that this intrinsic flexibility is maintained in the context of the replicase/encapsidase dimer, providing even more degrees of conformational freedom, for example if one or both 627 domains, which are stabilized by few interactions with FluPol, dissociate from the surface. Experimental data will be necessary to verify the validity of both the assumptions inherent to this modelling and the true context of the very different interaction mechanisms of *hu*ANP32A and *ch*ANP32A revealed by NMR, but it already seems clear that a degree of convergence exists between the complementary information provided by NMR and cryo-EM.

In conclusion, by comparing the ternary complexes of *hu*ANP32A and *ch*ANP32A with NP, we identify two very different interaction mechanisms that provide the key to understanding the molecular basis of adaptive mutations in the C-terminal region of PB2 that are essential for influenza replication in human cells. While *ch*ANP32A can bind FluPol and NP simultaneously via distinct linear motifs, the deletion in *hu*ANP32A blocks this mode of colocalization in *hu*ANP32A. In order to function, IAV therefore must mutate residues on the surface of 627, completing and enhancing the basic interaction surface, allowing *hu*ANP32A to then simultaneously bind FluPol and NP via the same surface, thereby forming a multivalent ternary complex. The identification of a common, extended motif on the IDD of ANP32A that simultaneously binds both viral proteins hints at possible exploitation of this sequence for the development of potent peptide-based inhibitors of NP and FluPol.

## MATERIALS AND METHODS

### Constructs

Chicken ANP32a (*Gallus gallus*, XP_413932.3) was synthesised and codon optimised for expression in *E. coli* (GenScript, New Jersey, USA). Human ANP32a was over expressed using plasmid pQTEV-ANP32a, which was a gift from Konrad Büssow (Addgene plasmid 31563). Plasmids containing just the intrinsically disordered region of chicken (149-281) or human ANP32a (144-249) were cloned into a pET9a derived plasmid with an N-terminal His-tag and a TEV cleavage site (MGHHHHHHDYDIPTTENLYFQG).

A codon-optimised construct containing the 627-NLS domains of the PB2 subunit was synthesised (538-759) from avian H5N1 A/duck/Shantou/4610/2003 (Geneart, Regensburg, Germany) for expression in *E. coli*.^11^ A construct containing just the 627 domain (538-693) was cloned into a pET9a plasmid with an N-terminal His-tag and a TEV cleavage site (see above).^32^

The NP gene from strain A/WSN/33 (H1N1) with a His-tag at its C-terminal was cloned in the pET22 plasmid (Novagen). Single point mutation R416A was introduced through the Quick-Change kit (Stratagene) in order to induce monomerisation of the protein.^48^ The NP H17N10 gene from little yellow-shouldered bat/Guatemala/060/2010(H17N10) was cloned in the pETM11 plasmid with an N-terminal His-tag and a SUMO tag containing a TEV cleavage site. The single point mutation R415A was introduced for monomerization.^48^

### Protein expression and purification

All proteins were expressed in *E. coli* - BL21 (DE3) Novagen) or AI (Invitrogen). Cultures were grown in LB or M9 media and induced with IPTG for 16 h at 18 °C. Bacteria were harvested by centrifugation and resuspended in buffer A containing Tris-HCl pH 7.5 and 200 mM NaCl (buffer A for NP contained 1 M NaCl). Bacterial lysis was performed by sonication. All proteins were purified by affinity chromatography on Ni-NTA agarose (Qiagen), followed by incubation with TEV protease at 4 °C overnight coupled with dialysis into buffer A.

For NP over-expression, cells were grown in LB or M9 media until the OD600 reaches 0.8-1.0, follow by induction with 200 mM IPTG and overexpression for 16 h at 25 °C. Bacteria were harvested and resuspended in a lysis buffer containing Tris-HCl pH7.4, 300 mM NaCl, 2 mM MgCl2, 15 mM imidazole and RNase A. Bacterial cells were lysed by sonication. Clarified bacterial lysate was then incubated with Ni-NTA agarose (Qiagen) at 4 °C for 30 minutes. Ni-NTA beads were then washed with 10 column volumes of high salt buffer containing Tris-HCl pH 7.4, 1 M NaCl, 2 mM MgCl_2_ and 1 mM β-ME. Protein was then eluted with 300 mM Imidazole. The eluent was then dialyzed into Tris-HCl pH 7.4, 300 mM NaCl and 0.5 mM β-ME, then loaded onto a HiTrap Heparin HP affinity column (Cytiva life sciences). A salt gradient from 300 mM NaCl to 1 M NaCl was used during elution.

For PRE experiments, a single cysteine mutant at position 631 of the 627 domain was tagged using 4-maleimido-TEMPO. Briefly, purified cysteine mutants were reduced with an excess of DTT at 4 °C for 12 hrs, and then dialysed into 50 mM phosphate buffer pH 7.0 with 150 mM NaCl and no DTT. Afterwards, a 5-fold molar excess of 4-maleimido-TEMPO dissolved in DMSO was added to the reduced protein. The reaction was incubated for 12 hrs at 4 °C and then excess of TEMPO was eliminated through size exclusion chromatography.

### Isothermal titration calorimetry

Isothermal titration calorimetry was measured using MicroCal iTC200 (GE healthcare) at 25 °C. Experiments were performed typically by adding 2.5 μL of aliquots of 200 μM of human or chicken ANP32A into the microcalorimeter cell filled with 20 μM of monomeric H17N10 NP; the titration mixture was stirred at 750 rpm. Data were fitted using Origin 7.0 with the ITC plugin from MicroCal.

### Nuclear magnetic resonance experiments

All NMR experiments were acquired, unless indicated, in 50 mM Tris-HCl buffer pH 6.5, 50 mM NaCl and 10% D_2_O on Bruker spectrometers with 1H frequencies of 600, 700, 850 and 950 MHz equipped with cryoprobes operating at 20 °C. All spectra were processed using NMRpipe^66^ and analysed in NMRFAM Sparky.^67^

For the RNA competition experiments, TROSY spectra of 1:1 mixtures of 25 μM of ^15^N *hu*ANP32A with NP were measured in the absence and presence of a 16-mer poly-UC oligonucleotide RNA repeat at a 2, 4 or 8 fold excess at 700 MHz.

^15^N R_1ρ_ relaxation rates were measured at 20 °C using a spin lock of 1.5 kHz.^68^ A typical set of relaxation delays included points measured at 1, 15, 30, 50, 70, 120 and 230 ms, including repetition of one delay. Relaxation rates were extracted using in-house software and errors were estimated using a noise-based Monte Carlo simulation. Transverse cross-correlated ^15^N-^1^H CSA/DD cross relaxation rates (*η_xy_*) were measured as previously described.^69^

Paramagnetic relaxation enhancement effects used to probe the formation of the ternary complex were measured from peak intensity ratios of ^15^N-TROSY spectra between a paramagnetic and a diamagnetic sample which was incubated with 2 mM ascorbic acid. The samples contained 100 μM ^15^N-labelled NP (H1N1, 416A) and TEMPO-labelled 627(K), with or without huANP32A (1:1:0 or 1:1:1).

^15^N CPMG relaxation dispersion experiments^70–72^ were acquired at 20 °C using a constant time relaxation delay of 32 ms, a ^1^H decoupling field of 11 kHz and 14 CPMG frequencies (ν_CPMG_) ranging from 31.25 to 1000 Hz, including repeats for estimation of uncertainties on peak intensity. Uncertainties on R_2eff_ values were propagated from the peak intensity uncertainty using a Monte Carlo approach. All relaxation dispersion data was analysed using the program ChemEx (https://github.com/gbouvignies/ChemEx).

### Modelling of ternary complexes

The recently published complex of ICV FluPol asymmetric dimer,^37^ bound to *hu*ANP32A was used as a template to construct a putative model of an equivalent IAV complex by superposing equivalent domains and modeling the replicase in elongation state, with the 5’ end of the product extending into the encapsidase 5’ hook binding site.

Ensemble descriptions of the ternary complex of *hu*ANP32A:NP:627(K)NLS, accounting for the disordered domain of *hu*ANP32A and *ch*ANP32A, were constructed using the flexible-meccano algorithm,^50,51^ which samples conformational space by randomly sampling free-energy surfaces describing the backbone dihedral angles ϕ/ψ for each residue in the disordered domain. Steric hindrance due to the presence of the FluPol domains, NP and the bound RNA are accounted for using hard sphere repulsion. Reasonable constraints were used to restrict conformational space to test certain selected scenarios. Namely – (a) The IDD of the binding region of *hu*ANP32A (170-221) should be within 15Å of the surface residues of 627 (FluPol_E_ or FluPol_R_) that are shifted upon interaction and (b) residues from this region should lie within 40Å of the RNA exit channel. This distance was chosen as representative of the distance between the mid-point of a 24 nucleotide RNA strand, and its two tethered ends (assumed immobilized on FluPol_R_ and FluPol_E_). A copy of NP was also placed in the RNA exit groove, with its RNA binding surface at 40Å of the RNA exit channel and the sampling repeated, this time with the constraint that the binding region of *hu*ANP32A should lie within 15 of the RNA binding grooves, where ANP32A is also assumed to bind. For the *ch*ANP32A the FluPol binding site should be within 15Å of the surface residues of either 627 domain, and the NP binding site within 40Å of the RNA exit channel. Different conformers that fulfil these restraints are shown in the figures to illustrate feasibility.

### Cells

HEK-293T ANP32AB knock-out (KO) cells were kindly provided by M Budt and T. Wolff (Robert Koch Institute, Berlin, Germany) and were grown in complete Dulbecco’s modified Eagle’s medium (DMEM, Gibco) supplemented with 10% fetal bovine serum (FBS) and 1% penicillin-streptomycin.

### Plasmids for cellular assays

The A/WSN/33 (WSN, H1N1) pcDNA3.1-PB2, -PB1, -PA and pCI-NP plasmids were described previously.^73,74^ The pFluPol-HA-mCherry plasmid was generated by replacing the internal sequence of the pFluPol-HA plasmid of the WSN reverse genetic system^75^ in between 125 nucleotides from the 5’end and 77 nucleotides from the 3’end by mCherry through standard PCR cloning followed by deletion of ATGs in the HA open-reading frame by an adapted site-directed mutagenesis protocol as described previously.^74^ The coding sequence of huANP32 was generated by gene synthesis (GenScript Biotech) and cloned into the pCI vector by standard PCR cloning. The indicated mutations were introduced by site-directed mutagenesis.

### vRNP reconstitution assay

The day before transfection 3.10^4^ HEK-293T ANP32AB KO cells were seeded in 96-well transparent plates (Corning 3599). The next day, cells were co-transfected using PEI (PEI MAX MW 40,000, Polysciences) with the following plasmids: 25 ng each of pcDNA3.1 PB2, PB1, PA, 50 ng pCI-NP, 50 ng pFluPol-HA-mCherry, and 50 ng pCI-empty or huANP32A (WT or the indicated mutant). Fluorescence signals were acquired using the Incucyte S3 (Essen Bioscience) every 4 h (10x objective, 5 fields per well) and were analysed using the Incucyte Live-Cell Analysis System (Sartorius) and are represented as RCU x µm²/Well.

## Funding

This work was supported by the European Research Council Advanced Grant DynamicAssemblies under the European Union’s Horizon 2020 research and innovation program (grant agreement number 835161) (M. B.).

The work used the platforms of the Grenoble Instruct-ERIC centre (ISBG; UMS 3518 CNRS-CEA-UJA-EMBL) with support from FRISBI (ANR-10-INSB-05-02) and GRAL (ANR-10-LABX-49-01) within the Grenoble Partnership for Structural Biology (PSB).

ACZ received funding from the European Union’s Horizon 2020 research and innovation programme under the Marie Skłodowska-Curie grant agreement No. 796490 and HFSP postdoctoral HFSP fellowship LT001544/2017.

Financial support from the IR-RMN-THC Fr3050 CNRS for conducting the research is gratefully acknowledged.

IBS acknowledges integration into the Interdisciplinary Research Institute of Grenoble (IRIG, CEA).

We would like to thank M. Budt (Robert Koch Institute, Berlin, Germany) for providing HEK-293T ANP32AB knock-out (KO) cells.

## Supporting Information

Supporting information is available for this paper, including full presentation of the relaxation dispersion curves of NP titrated into solutions of *hu*ANP32A, cross correlated cross relaxation of different admixtures of 627(K)-NLS: *hu*ANP32A admixtures and ITC of human and avian ANP32A with NP.

## Supporting Information

**Figure S1.**
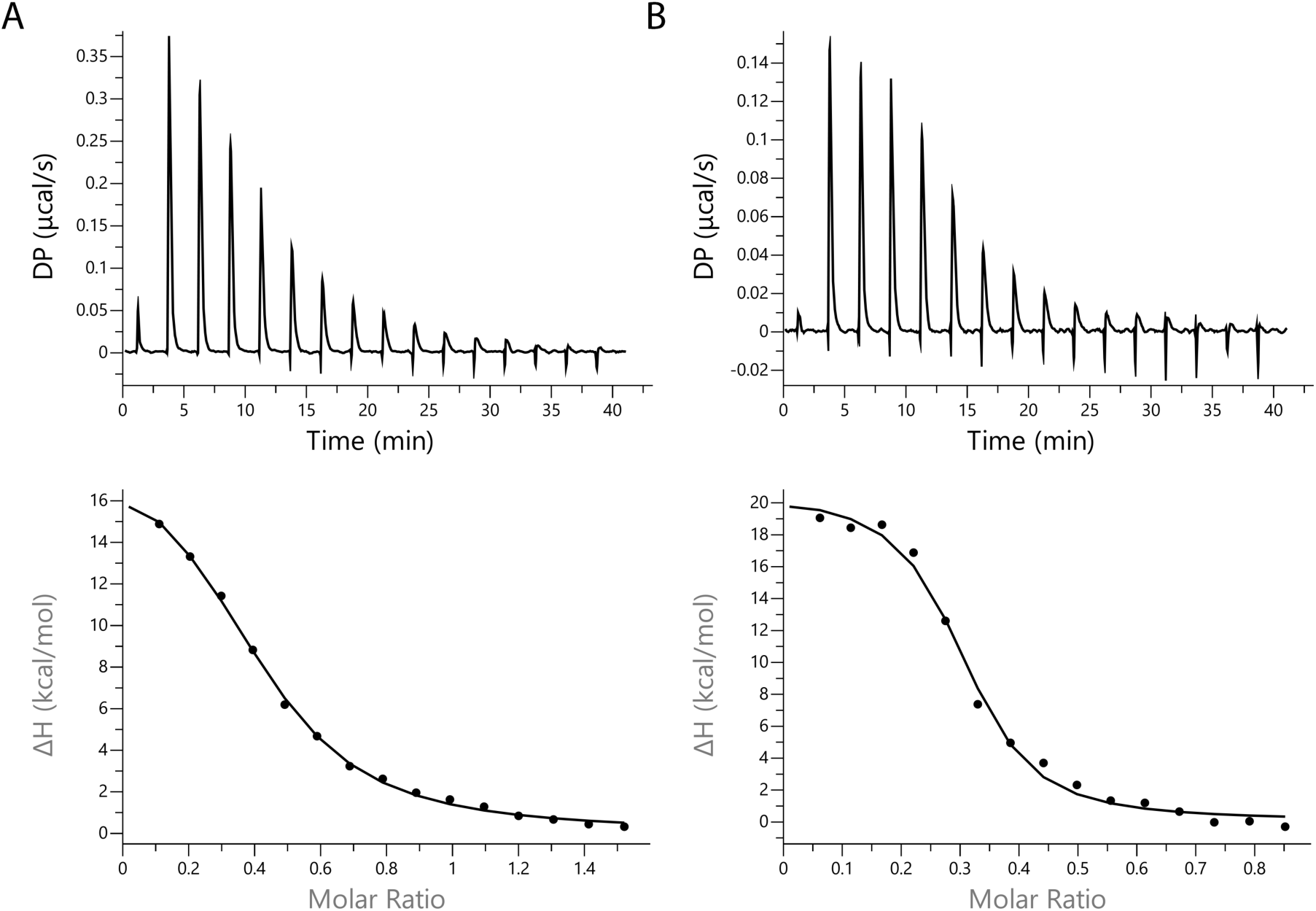
ITC shows higher affinity of NP for *av*ANP32A than *hu*ANP32A. **A -** ITC curve of *hu*ANP32A for monomeric NP; 150 μM of huANP32A were titrated into a cell with 20 μM of NP. Data were fitted to a simple bimolecular binding model, resulting in fitted parameters: ΔH= 19.1±0.75 kcal.mol^-1^, TΔS=26.9 kcal.mol^-1^, K_D_=1.7±0.2 10^-6^M, N=0.40±0.01. **B** - ITC curve of avANP32A for monomeric NP; 42 μM of avANP32A were titrated into 10 μM of NP. Data were fitted to a simple bimolecular binding model, resulting in fitted parameters: ΔH= 20.5±0.9 kcal.mol^-1^, TΔS=30.0 kcal.mol^-1^, K_D_=0.126±0.32 10^-6^M, N=0.29±0.01.

**Figure S2.**
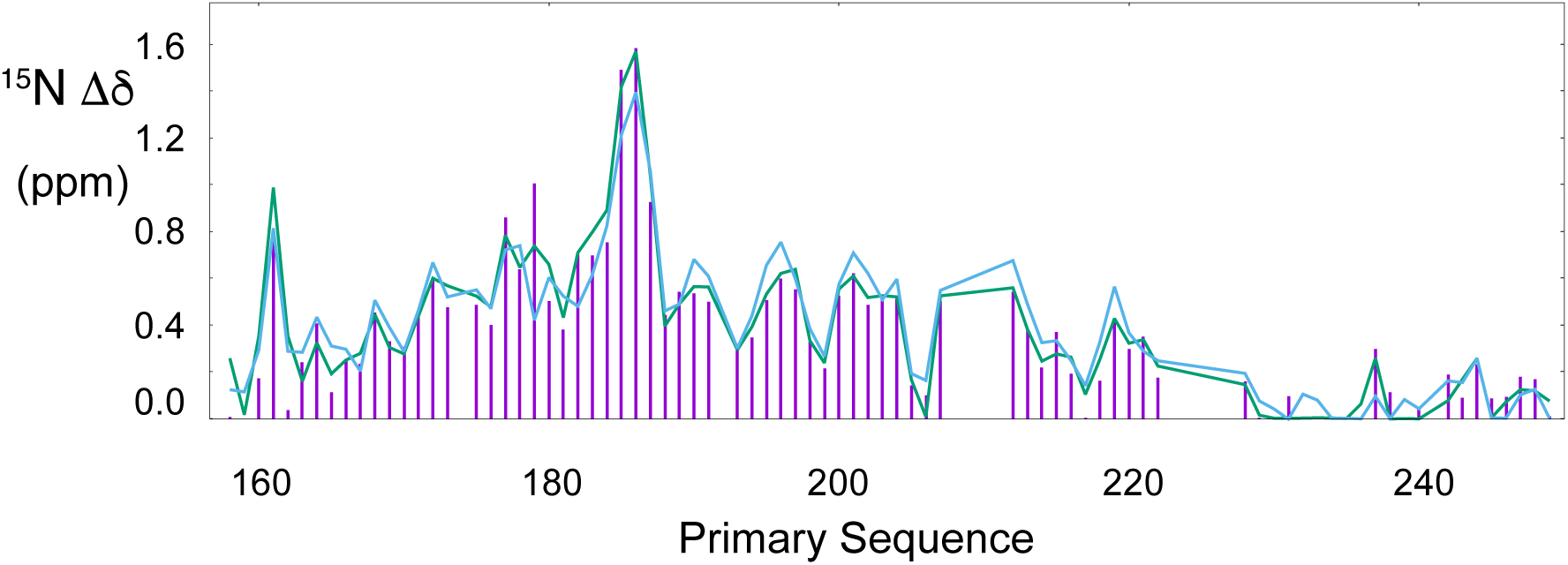
Chemical shift differences between free and bound ANP32A. Analysis of CPMG relaxation dispersion data at each independent admixture using a single 2-state exchange model with common exchange rates and populations throughout the protein, results in essentially identical chemical shift perturbations between the free and bound states at each admixture (purple – 5% admixture, blue – 10% admixture, green – 20% admixture). This testifies to both the consistent nature of the experimental data, and the applicability of the 2-state binding model. Populations of the bound states were determined to be 0.037±0.005, 0.067±0.007 and 0.117± 0.009 respectively.

**Figure S3.**
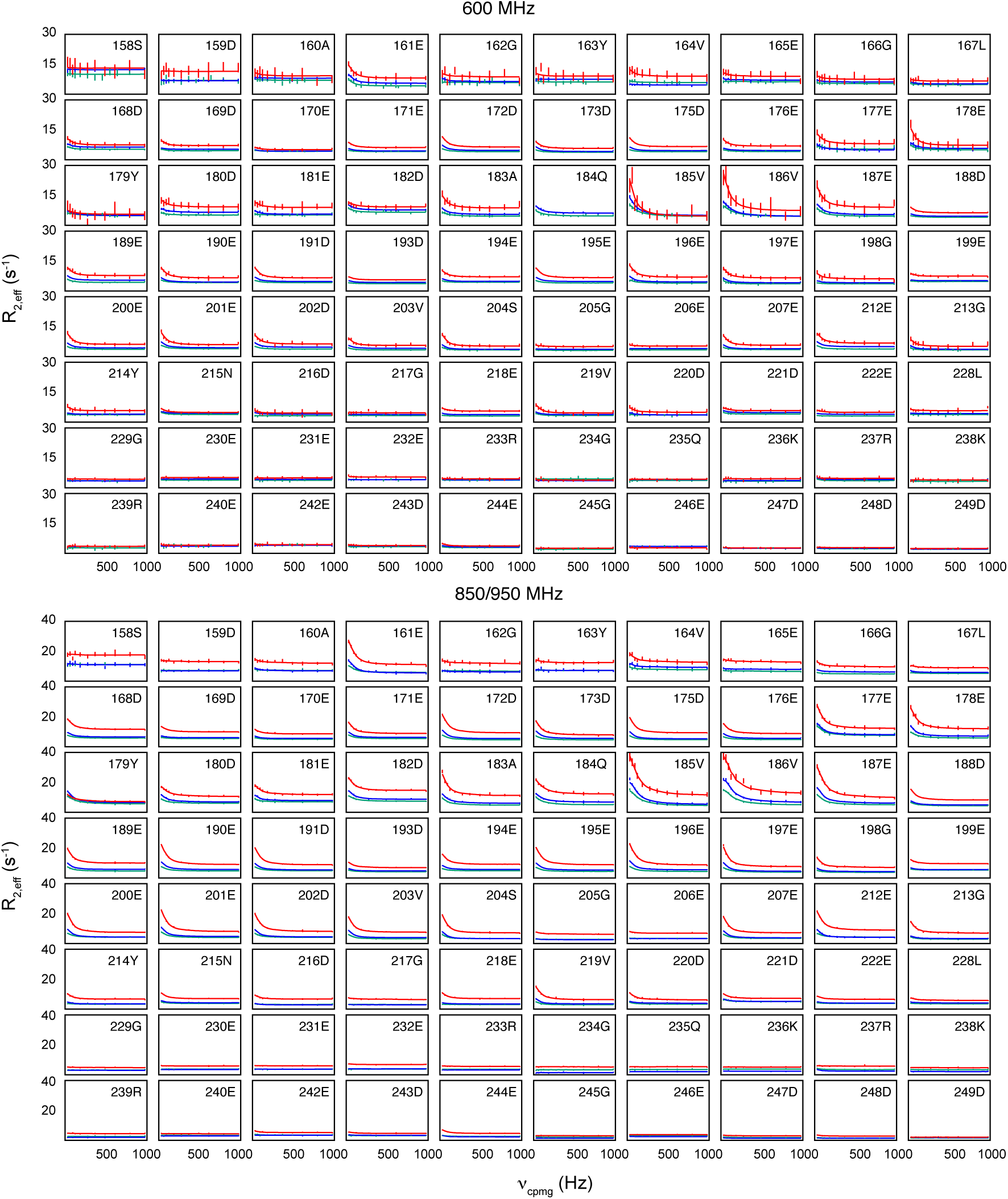
^15^N CPMG Relaxation dispersion measured in *hu*ANP32A IDD in complex with NP. ^15^N CPMG relaxation dispersion was measured at two magnetic field strengths (600MHz and 850 or 950MHz) on *hu*ANP32A samples containing three different admixtures of NP (green - 5%, blue - 10% and red - 20%). Data from each admixture were fitted to a single two-site exchange process (fitted values are shown as lines). On the basis of fitted populations, the effective K_D_ is 50.5±12.2μM. Exchange rates for 5%, 10% and 20% admixtures are closely clustered (302±25, 249±62 and 291±31) suggesting that exchange rate is dominated by k_off_.

**Figure S4.**
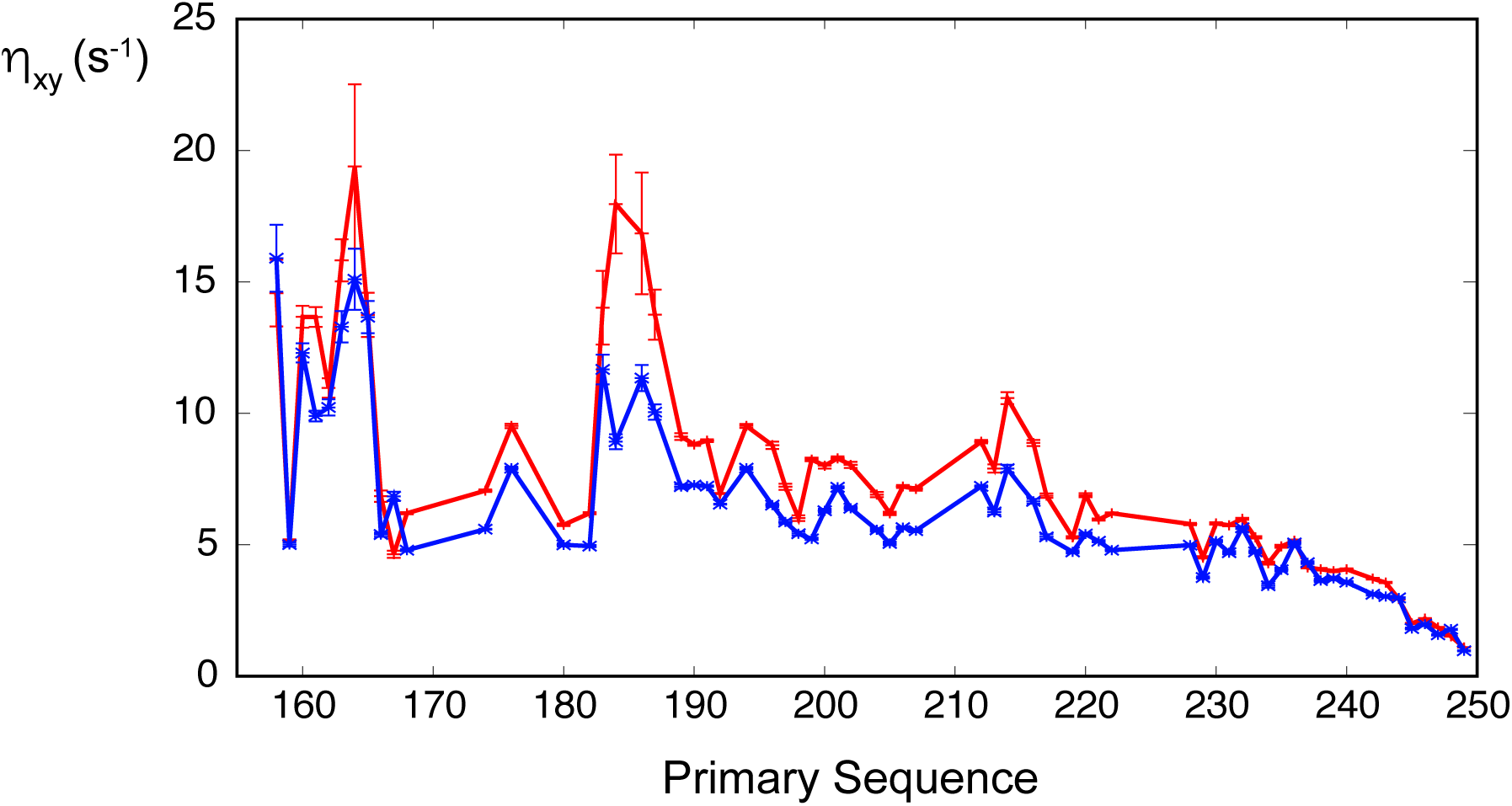
Cross-correlated dipole-dipole/CSA cross relaxation of binary and ternary complexes. (Blue) Cross-correlated dipole-dipole/CSA cross relaxation was measured at 850MHz on *hu*ANP32A bound to 627(K) (1:2 admixture), compared to (red) cross-correlated dipole-dipole/CSA cross relaxation measured at 850MHz on ANP32A bound to 627(K) and NP (1:2:0.1 admixture). The same phenomenon, of increased transverse relaxation in the ternary mixture compared to the binary mixture, seen in the rotating frame relaxation experiments (figure 4A), is seen here, ruling out a chemical shift exchange effect.

